# Structural Basis for C^8^ methylation of 23S ribosomal RNA by Cfr

**DOI:** 10.64898/2026.02.27.707579

**Authors:** Olga A. Esakova, James Jung, Hyunwook Lee, Sung Hyun Cho, John Alumasa, Erica L. Schwalm, Tyler L. Grove, Matthew R. Bauerle, Susan L. Hafenstein, Zhiheng Yu, Squire J. Booker

**Affiliations:** The Department of Chemistry, The Pennsylvania State University, University Park, Pennsylvania 16802, USA; the Department of Biochemistry and Molecular Biology, The Pennsylvania State University, University Park, Pennsylvania 16802, USA; The Huck Institutes of the Life Sciences, The Pennsylvania State University, University Park, Pennsylvania 16802, USA; the Howard Hughes Medical Institute, The Pennsylvania State University, University Park, Pennsylvania 16802, USA; The Department of Chemistry (School of Arts and Sciences), University of Pennsylvania, Philadelphia, PA 19104, USA; the Department of Biochemistry and Biophysics (Perelman School of Medicine), University of Pennsylvania, Philadelphia, PA 19104, USA; the Howard Hughes Medical Institute, University of Pennsylvania, Philadelphia, PA 19104, USA; Janelia Research Campus, Howard Hughes Medical Institute, Ashburn, VA 20147, USA; Department of Chemistry and Biochemistry, Miami University of Ohio, Oxford, OH 45056; Department of Biochemistry, Albert Einstein College of Medicine, Bronx, New York 10461; The Hormel Institute, University of Minnesota, Austin, MN 55912; College of Biological Sciences, University of Minnesota, Twin Cities, St. Paul, MN 55108; Mayo Clinic, Department of Infectious Diseases, Rochester, MN 55905

## Abstract

Cfr methylates C^8^ of adenosine 2503 (A2503) in 23*S* ribosomal RNA (rRNA) and will also methylate C^2^ of A2503 after methylating C^8^. C^8^ methylation confers resistance to more than five classes of clinically used antibiotics, highlighting it as a worrisome mechanism of antibiotic resistance. Here, we report the structure of Cfr, determined by cryogenic electron microscopy (Cryo-EM). Despite its small size (∼36 kDa), we exploit a transient protein–RNA crosslink that forms during catalysis, which requires Cys105 to resolve. Using a Cfr Cys105Ala variant and an 87-nucleotide strand of rRNA, we isolate the crosslinked species and determine its structure to 3.0 Å resolution. Notably, the 87-mer rRNA adopts an L-shaped conformation characteristic of tRNAs, rather than the conformation it assumes in the ribosome.

**One Sentence Summary:** Cryo-EM structure of Cfr, a radical S-adenosylmethionine methylase that confers antibiotic resistance

## Introduction

Infections caused by antibiotic-resistant bacteria remain a significant threat to human health worldwide. In the United States, about 2.8 million infections occur annually, resulting in over 35,000 deaths^1^. Over 700,000 people, including 214,000 newborn babies, die annually from such infections^1, 2^. Multidrug-resistant strains are primarily responsible for these sobering statistics^3^. The prevalence of such strains has led to the emergence of fast-spreading ‘superbugs’ unresponsive to current treatment regimens^4, 5^. These pathogens employ myriad strategies to resist antibiotics, including modifying macromolecules that antibiotics target within bacterial cells^6, 7^. One such strategy is methylating adenosine 2503 (A2503) of 23*S* ribosomal RNA (rRNA) by the product of the *cfr* gene, which confers resistance to several classes of clinically important antibiotics, including phenicols, lincosamides, oxazolidinones (including linezolid), pleuromutilins, streptogramin A, hygromycin A, and the 16-membered macrolides josamycin and spiramycin ^8-12^. Recent studies of Cfr-modified ribosomes have provided a more precise understanding of how resistance to these antibiotics arises. In some cases, the modified nucleotide sterically impedes antibiotic binding. In other cases, it results in a reorganization of nucleotides in the antibiotic binding pocket ^10, 11^.

Cfr is highly related to RlmN, which also methylates A2503 of 23*S* rRNA but at C^2 13, 14^. However, unlike Cfr, RlmN will also methylate C^2^ of adenosine 37 (A37) in several transfer RNAs (tRNA)^15^. Structural studies of RlmN^16, 17^ and mechanistic studies of RlmN and Cfr unveiled an unprecedented radical-dependent methylation mechanism shared by the two enzymes^14, 16, 18-23^. Both are members of the radical *S*-adenosylmethionine (SAM) superfamily of enzymes, which use a [Fe_4_S_4_] cluster to cleave SAM reductively to afford a 5’-deoxyadenosyl 5’-radical (5’-dA•)^24-28^. In RlmN’s and Cfr’s first catalytic step, a methyl group from SAM is transferred to a conserved Cys residue of the protein via an S_N_2 mechanism. After *S*-adenosylhomocysteine (SAH) departs, the binding and reductive cleavage of a second SAM molecule affords the 5’-dA•, which abstracts a hydrogen atom (H•) from the *S*-methylcysteine (mCys) residue. The resulting methylene radical adds to the substrate’s C^2^ (RlmN) or C^8^ (Cfr) atom, forming a protein–RNA crosslink with an unpaired electron. A second Cys (C105 for Cfr and C118 for RlmN) abstracts the C^2^ (RlmN) or C^8^ (Cfr) proton, promoting the breakdown of the crosslinked species through a radical fragmentation of the C-S bond and generating a thiyl radical and an enamine. Catalysis is complete upon reduction of the thiyl radical and tautomerization of the enamine to the methylated product upon return of the abstracted proton (**Figure S1**)^17, 18, 21^.

Several X-ray crystal structures of RlmN have been determined, including a structure of the crosslinked species with tRNA^Glu16, 17^. No structures of an active, wild-type (WT) Cfr protein have been forthcoming despite more than a decade of attempts to obtain diffraction-quality crystals. Moreover, no structures of RlmN have been determined with a fragment of 23*S* rRNA. Herein, we report the Cryo-EM structure of Cfr crosslinked to an 87-nucleotide (87-mer) 23*S* rRNA fragment that incorporates A2503. We find that the bound 87-mer rRNA fragment mimics the L-shape structure of the tRNA in the structure of RlmN crosslinked with tRNA^Glu^. The structure reveals that most interactions between Cfr and the RNA substrate occur with the phosphate and sugar moieties of the nucleic acid, enabling it to recognize shape rather than a specific sequence. Near the active site of Cfr, the RNA backbone is distorted, allowing A2503 to access the enzyme’s catalytic machinery.

## Results

Cryo-EM is most often used to determine the atomic-level structures of macromolecules or macromolecular assemblies larger than 100 kDa^29, 30^. Major advances in electron microscopy have enabled the successful reconstruction of high-resolution density maps for macromolecules with large molecular weights using single-particle analysis (SPA)^29, 31^. The size constraint results from the low signal-to-noise ratio generated by smaller proteins and the difficulty in aligning the smaller single particles^30^. Because the molecular weight of *Staphylococcus aureus* Cfr (is only ∼40 kDa, we leveraged the size of its macromolecular substrate by engineering a variant that generates a covalent Cfr-rRNA complex during catalysis to address the size requirement. Because Cfr requires C105 to initiate the breakdown of its covalent complex with its rRNA substrate, C105A or C105S substitutions form a crosslinked species that can be purified and characterized^18, 21^.

We identified an 87-mer fragment (25.4 kDa) of 23*S* rRNA that supports robust Cfr activity and was sufficient for studying the Cfr-rRNA complex by SPA (**Figure 1**). This rRNA fragment spans nucleotides 2496-2582 and includes stems 90, 91, and 92 (*E. coli* numbering; see Materials and Methods for details). Although Cfr is oxygen-sensitive, the rRNA fragment appears to provide sufficient protection to allow grids to be prepared aerobically with less than 1 min of exposure to air. Graphene monolayer grids, fabricated via chemical vapor deposition (CVD), enabled the formation of extremely thin ice, achieving over 95% transmittance of the incident beam dose rate. Moreover, the CVD-prepared graphene monolayer efficiently suppressed beam-induced motion to a sub-0.5-1 Å level on 2.0 μm diameter holes (**Figure S2E**), comparable to HexAuFoil grids with 300 nm holes. Sub-1 Å-level beam-induced motion eliminated the need for particle polishing^32^. Adsorption of Cfr particles to the uniform CVD-prepared graphene monolayer surface enabled more consistent particle Z-positions, resulting in a more accurate particle contrast transfer function (CTF) estimation, which is crucial for determining the structure of small molecular weight targets by Cryo-EM^33^. Adsorption to the graphene monolayer surface protected the small Cfr particles from the otherwise detrimental effects at the air-water interface.

**Figure 1.**
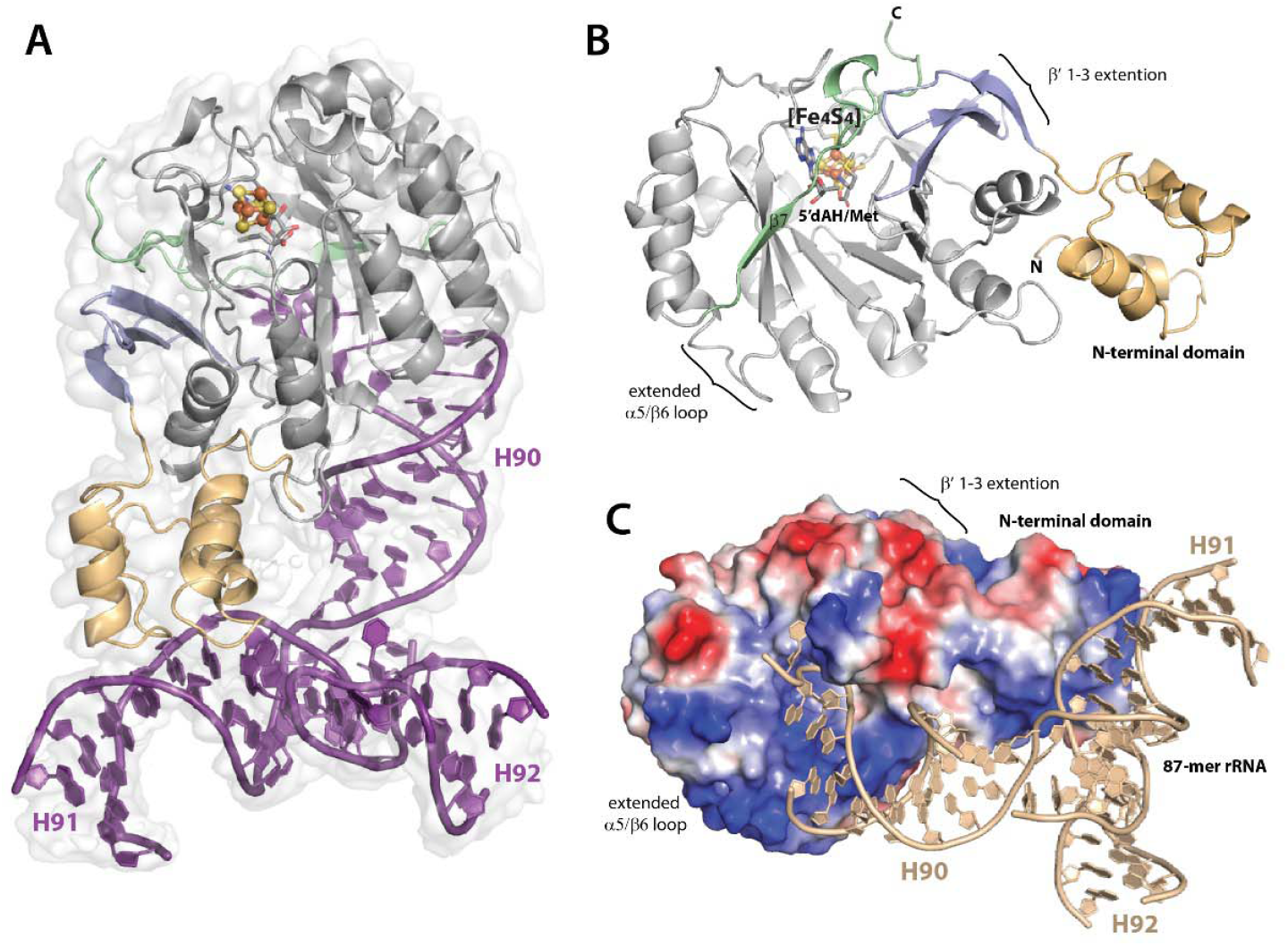
A cartoon representation of (**A**) cross-linked complex of CfrC105A/87-mer complex overlayed on the Cryo-EM map; (**B**) Cfr domain architecture. Color code: N-terminal domain (tan); β-sheet extension (light blue); radical SAM domain (light grey); C-terminal (green). (**C**) The electrostatic surface potential of Cfr contoured at -70 mV (red, negative) and 70 mV (blue, positive), 87-mer rRNA is tan color.

### The overall Cryo-EM structure of Cfr crosslinked complex

The map for the Cfr C105A/87-mer complex was reconstructed at a near-atomic resolution of ∼3.0□Å, with an overall resolution of 3.02 Å and a local resolution ranging from 2.9 to ∼3.3 Å (**Table S1, Figure S2**). In this complex (**Figure 1A**), the structure of Cfr exhibits a high degree of similarity to that of *E. coli* RlmN, with an RMSD of 1.1 Å for 229 Cα-atoms (**Figure S3**). The overall architecture of the N-terminal domain of Cfr is based on the HhH2 fold, typically observed in non-sequence-specific DNA and RNA binding proteins, and consists of four short α-helices (**Figure 1B**). 3D variability analysis (3DVA) using cryoSPARC v4 (movie in SI) reveals a notable level of conformational flexibility in the N-terminal domain, with a loop spanning Phe59-Asn65 acting as a hinge region. This loop may have a limited attainable resolution using cryo-EM^34^. Like in RlmN, the N-terminal domain of Cfr contains a positively charged region that plays a crucial role in rRNA binding (**Figure 1C**).

The subsequent domain shares common features with all radical SAM (RS) enzymes^35-37^. It adopts a partial (βα)_6_ triosephosphate isomerase (TIM) barrel fold containing a conserved CX_3_CX_2_C motif that contributes ligands for binding the [Fe_4_S_4_] cluster. The TIM barrel fold is prolonged by a β7-strand extending into the C-terminal unstructured region. The N-terminal domain is connected to the RS domain through a β-sheet extension (β’1-3 strands) that aligns on one side with the first β-strand of the RS domain and supports the C-terminal unstructured region on the other side. A comparison of the RS domain architecture of Cfr and RlmN reveals that RlmN has an α1/β2 loop extension. By contrast, Cfr has a loop extension between α5 and β6. Specifically, the elongated α5/β6 loop, located near the substrate-binding region of the RS domain, carries a predominantly positive charge, highlighting its potential role in binding large rRNA substrates. Notably, the Cfr structure in the complex lacks a C-terminal helix, a distinctive feature of RlmN (**Figure S3**).

### The rRNA binding sites

Like RlmN, Cfr acts on ‘naked’ 23*S* rRNA^14^. The Cryo-EM structure of the Cfr/87-mer complex (**Figure 1**) reveals the presence of the crosslinked 87-mer rRNA fragment (**Figure S4A**), although only spanning nucleotides 2501 to 2582 (82 nucleotides). The reconstructed Cryo-EM density map supports most of the 87-mer’s sequence, which incorporates stems H90, H91, and H92. The map lacks densities for regions 2526-2537 of H91, 2551-2557 of H92, and 2571-2572 of H90. The secondary structure of the rRNA fragment in the complex (**Figure S4B**) mimics that of the rRNA in the ribosome (**Figure S4A**). However, it demonstrates more disordered density in some base-pairing regions that do not interact directly with Cfr. When overlaying the H90 stem-loop region of 23*S* rRNA from an assembled ribosome (PDB ID: 2AWB) with that of the 87-mer (**Figure S4C**), different orientations of H91 and H92 are observed. This observation suggests that the rRNA region incorporating H91 and H92 undergoes a ∼180° flip during ribosome maturation.

H92 is the most distant part of the rRNA in the crosslinked complex. It carries a highly conserved U^m^2552 modification, which is essential for selecting the correct tRNA at the ribosomal A site. U2552 is typically methylated at the 2’-OH group by the SAM-dependent enzyme RlmE, forming U^m^2552 on the assembled 50*S* ribosomal subunit. The lack of the U^m^2552 modification results in growth defects, reduced translation rates, incorrect readthrough of stop codons, and reduced stability of 70*S* ribosomes^38^. However, U2552 in the crosslinked complex is not involved in interactions with Cfr. The N-terminal domain of the enzyme is anchored to the central part of the 87-mer, from which the three stems (H90, H91, and H92) diverge. Tyr24 from the N-terminal domain participates in π-π stacking with A2565, located between H90 and H92. The side chains of Arg25 form hydrogen bonds (H-bonds) with the oxygen of the phosphate group of G2523 and the 3’-O of U2522, while the α-carbonyl oxygen of Val46 and the side chain of Gln28 form H-bonds with the 2’-OH group of U2522 (H91). The oxygen of the Lys45 backbone amide is in H-bonding distance to the 2’-OH group of G2523. The C_α_ nitrogen and the ε-amino group of Lys49 form H-bonds with the 5’-O and 3’-O of G2524 (H91), respectively. The nitrogen of the Gln36 side chain forms H-bonds with the 3’-O and the 2’-OH group of G2543 in H91 (**Figure 2A, B**). The N-terminal domain interacts primarily with the RNA backbone of the H91 stem, with nucleotides 2522-2554 forming one strand of hairpin H91 and G2543 as the only nucleotide from the opposite strand involved in the interaction. The structure of the Cfr/87-mer complex suggests that interactions between the N-terminal domain and RNA are not sequence-dependent; instead, they are likely directed toward recognizing the RNA tertiary structure. Notably, the N-terminal domain stabilizes the ribose-phosphate backbone near the turns between H90 and H91, H91 and H92, and H92 and H90.

**Figure 2.**
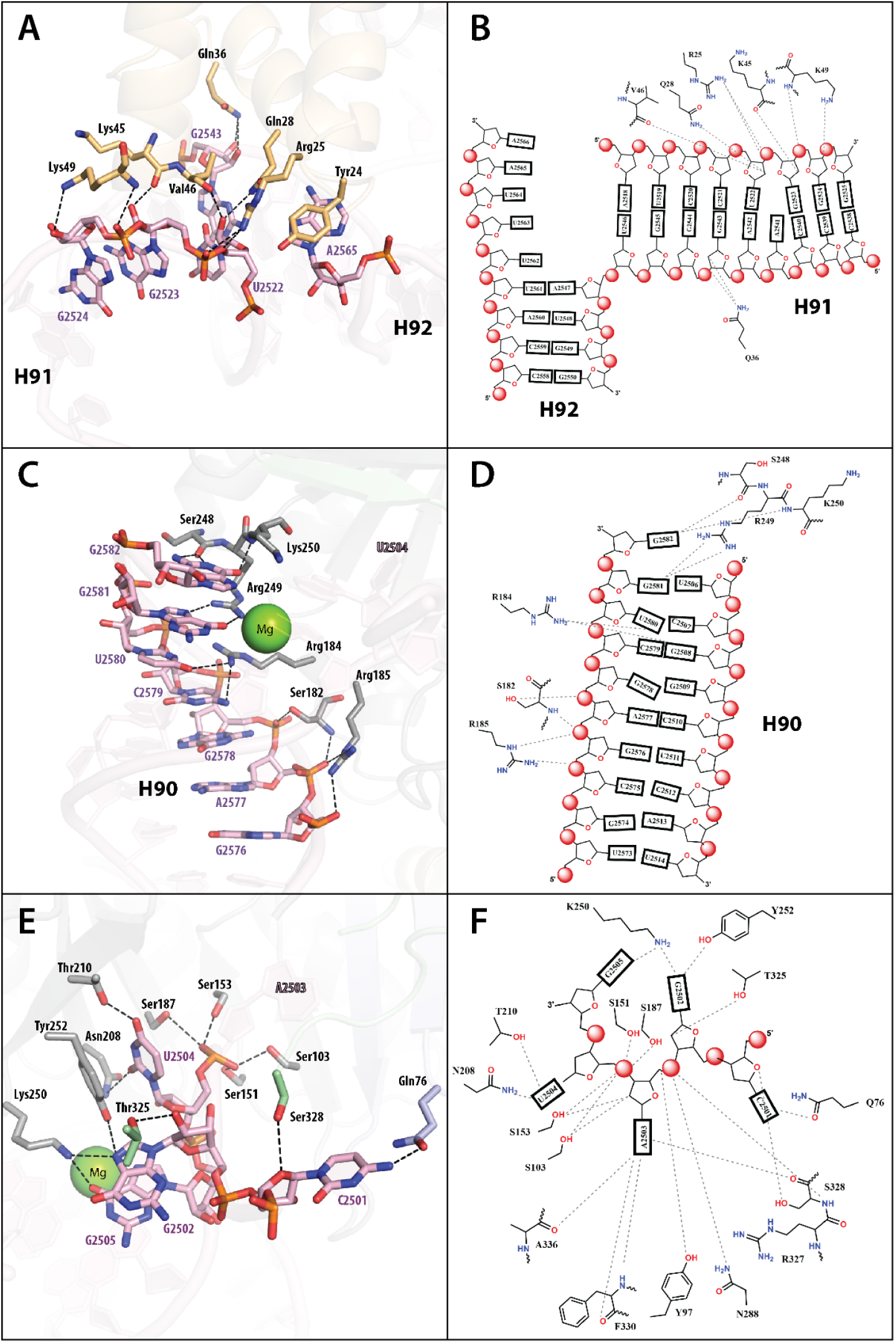
The representation of protein-RNA interaction in Cfr C105A/87-mer rRNA cross-linked complex (**A**) N-terminal domain of Cfr with H91 and H92 stems of rRNA and (**B**) scheme of H-bonds interactions; (**C**) RS-domain of Cfr with H90 stem of rRNA and (**D**) scheme of H-bonds interactions; (**E**) Cfr with single stranded region of rRNA (2500-2506), (**F**) scheme of H-bonds interactions. Color code: N-terminal domain (tan); β-sheet extension (light 10 blue); radical SAM domain (light grey); C-terminal (green).

The RS domain specifically coordinates the H90 stem (**Figures 2C and 2D**). The H90 stem in the ribosome structure is formed by nucleotides 2507 to 2517, which base-pair with nucleotides 2567-2582. However, in our structure, the H90 stem has a shorter base-pairing region (**Figures S4A and B**), and the RS domain interacts exclusively with only one of its strands (**Figures 2C** and **D)**. The positively charged region formed by amino acids in the RS domain binds to the H90 stem (**Figure 1C**). The amide nitrogen of Lys250 and the carbonyl oxygen of Ser248 H-bond with N^2^, N^1^, and O^6^ of G2582, respectively. Arg249 forms H-bonds with O^6^ and N^7^ of G2581. Arg184 is within H-bonding distance from two nucleotide bases, U2580 (O^4^) and C2579 (N^4^), and additionally forms a cation-π interaction with C2579. The OH and amide nitrogen of Ser182 coordinate the ribose-phosphate backbone of G2578 and A2577, respectively, and the guanidinium moiety of Arg185 coordinates the ribose-phosphate backbone of A2577 and G2576 (**Figure 2C, D**). Position 2580 in 23*S* rRNA is pseudouridine (□2580), which is installed by the pseudouridine synthase RluC^39^. Although not in our structure, this pseudouridine might be additionally stabilized by H-bonds with Arg184, given that the distance between C^5^ (i.e., N^1^ of pseudouridine) of U2580 and the guanidino group of Arg184 is only 4 Å.

Nucleotides 2501 to 2506 of the 87-mer form a single-stranded region. The 87-mer also incorporates nucleotide C2498, which is typically methylated on the 2’-OH by RlmM. Loss of RlmM function retards *E. coli* growth^40^. C2498 is a highly disordered nucleotide and was not modeled due to a lack of clear density. The first nucleotide supported by the density map in the single-stranded region is C2501. Its N^4^ atom forms H-bonds with the O of the Gln76 side chain and the 3’-O of the OH-group of Ser328 (**Figure 2E, F**). Additionally, C2501 is near an aromatic pocket formed by Tyr97, Trp101, and Phe330. Typically, C2501 is modified by RlhA, which installs an OH group at C^5^, forming 5-hydroxycytidine (ho^5^C2501). The ho^5^C2501 modification affects translation efficiencies and is beneficial for *E. coli* cells in adapting to oxidative stress^41^. Although the effect of this modification on Cfr recognition is unknown, the 3.4 Å distance between C^5^ of C2501 and the N of the Gln76 side chain suggests that this modification could stabilize the rRNA-Cfr complex.

Nucleotides 2502 to 2505 form a U-turn, also observed in the 23*S* rRNA of the ribosome structure (**Figure S3C**), and mimic the tRNA^Glu^ bound to RlmN (PDB ID: 5HR7) (**Figure S5**). This U-turn region of the rRNA inserts deep into Cfr, allowing A2503 to establish multiple interactions within the active site. The U-turn is stabilized by π-π stacking between G2502 and G2505, and Lys250 supports this interaction via H-bonds to the nucleobases (**Figure 2E, F**). Trp101 is inserted inside the U-turn and makes hydrophobic interactions with the carbons of the ribose-phosphate backbone of G2502, U2504, and G2505 (**Figure 2E, F**). The first position of the U-turn is G2502, which forms a tilted edge-to-face π-stacking interaction with His286. Also, N^7^ and the 2’-OH group form H-bonds with the OH groups of Tyr252 and Thr325, respectively.

A2503, the nucleotide undergoing methylation, is bound tightly within the active site (**Figure 3**). U2504 of the 87-mer is Ψ in 23*S* rRNA, also formed by RluC^42^. The absence of this Ψ was found to significantly increase the susceptibility of bacteria to peptidyl transferase inhibitors^42^. Both Ψ at 2504 and m^8^A at 2503 are post-transcriptional modifications that evolved as intrinsic mechanisms of antibiotic resistance^43^. The Ψ modification at position 2504 does not affect Cfr or RlmN activity, but its replacement with adenosine completely abolishes RlmN activity^44^. The crosslinked structure of Cfr with the 87-mer reveals that the phosphate oxygen of U2504 is stabilized by four H-bonds with the side chains of Ser103, Ser151, Ser153, and Ser187. Additionally, the U2504 nucleobase shows π-π stacking with Tyr252; its O^4^ forms an H-bond with the OH-group of Thr210, and its O^2^ forms an H-bond with the side chain of Asn208 (**Figure 2E, F**). Overall, the single-stranded region from 2501 to 2505 exhibits strong interactions with the enzyme, involving nearly all the nucleotides. This region continues into the double-stranded region of the H90 stem, and this transition is coordinated by a Mg^2+^ ion.

**Figure 3.**
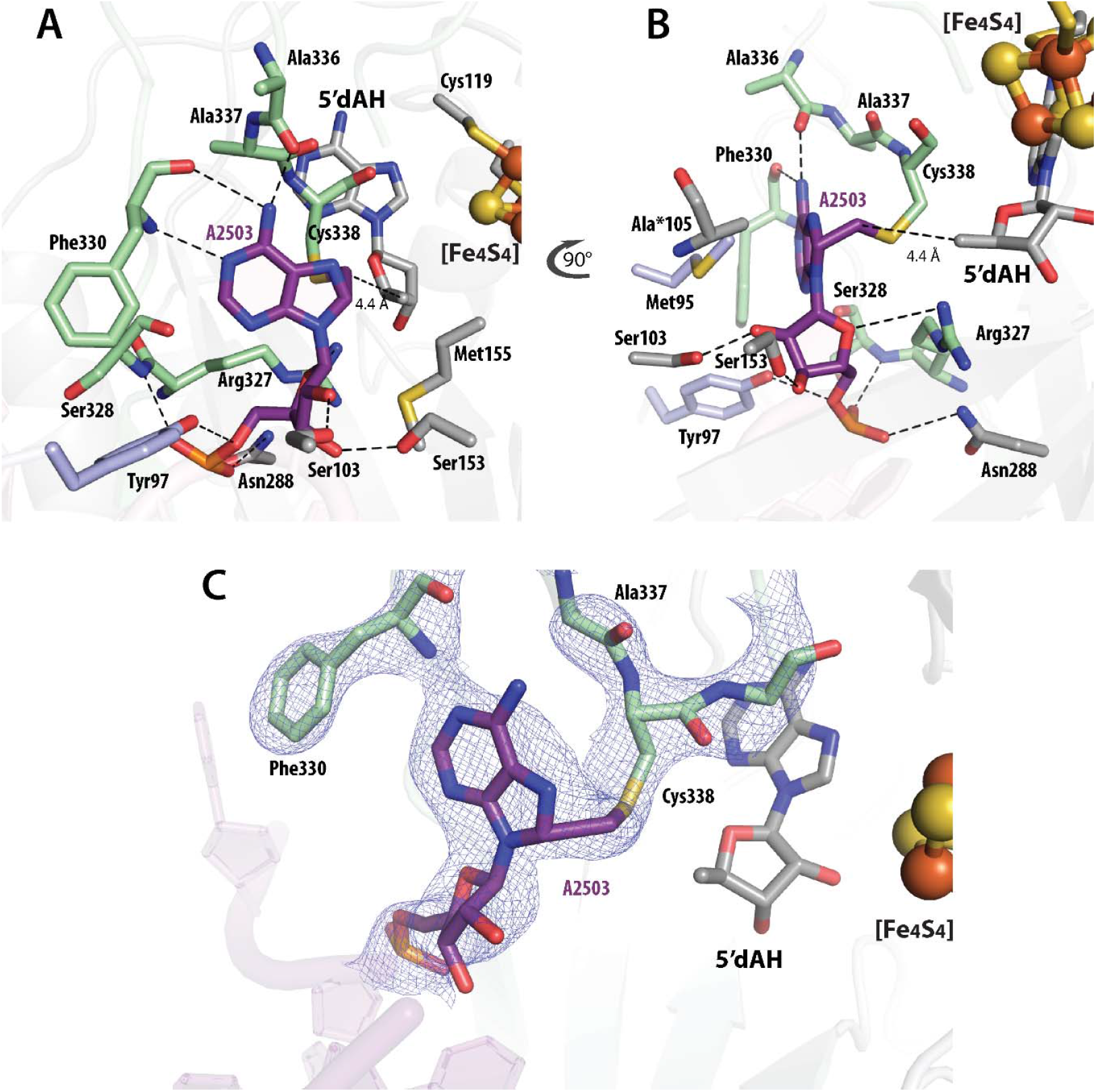
A cartoon representation of the active site of Cfr C105A/87-mer rRNA cross-linked complex (**A, B**). (**C**) EM density of the cross-linked Cfr C105A/87-mer rRNA intermediate. Color code of residues: β-sheet extension (light blue); RS-domain (light grey); C-terminal (green); A2503 of rRNA (purple).

### The Cfr active site

The structure of the Cfr C105A active site reveals the presence of 5′-deoxyadenosine (5′dAH) and L-methionine (Met), suggesting, as was found in the RlmN C118A tRNA^Glu^ crosslink structure, that 5’-dAH and Met are released after the crosslink is resolved and the methylated RNA leaves (**Figure S1**). Moreover, the Cfr complex exhibits an extended density at Cys338, which is related to the appended methylene group attached to C^8^ of A2503. The methylene carbon is within 4.4 Å of C5′ of 5’-dAH, appropriate for H• abstraction from the Cys-appended methyl group by the 5’-dA• (**Figure 3**). Cys105 abstracts the proton from the sp^3^-hybridized C^8^ of the crosslinked intermediate. However, this Cys is replaced by Ala to stabilize the protein–RNA crosslink by preventing its decomposition. Ala105 points in the direction of C^8^, and the distance between this carbon and C^β^ of Ala105 is 3.6 Å (**Figure 3B**). Modeling a Cys instead of Ala105 in the crosslinked structure shows a distance of ∼3.0 Å between the sulfur atom of Cys105 and C^8^ of A2503, suitable for proton abstraction. The proposed thiyl radical, generated on Cys338 after cleavage of the C-S bond, is believed to be reduced by electron transfer from the cluster, which is about 9.7 Å away.

Analysis of the interaction between A2503 and residues in the active site of Cfr suggests that the enzyme exhibits high specificity for this residue (**Figure 3**). The β’1-3 strand extension region stabilizes A2503 through van-der-Waals interactions with Met95 and H-bonds between Tyr97 and the 5’-O of the ribose. Ser103, Ser153, and Asn288 within the RS domain form H-bonds with the 3’-O and 2’-OH group of the ribose and the phosphate oxygen of A2503, respectively. The C-terminal unstructured region contributes H-bonds between the nitrogen and oxygen of Phe330 with N^1^ and N^6^ of A2503, the nitrogen of Ser328 with the phosphate oxygen, and Arg327 with the 4’-O of the ribose of A2503. The carbonyl oxygen of Ala336 forms an H-bond with N^6^ of A2503.

## Discussion

The structures of Cfr crosslinked to an rRNA fragment (determined by Cryo-EM) and RlmN crosslinked to tRNA^Glu^ (determined by X-ray crystallography) are remarkably similar, exhibiting an RMSD of 1.1 Å over 229 Cα-atoms. Although the shape and sequence of the rRNA substrate recognized by both enzymes are identical, only RlmN exhibits dual specificity, capable of installing a methyl group at C^2^ of A2503 and C^2^ of A37 in several tRNAs^15, 17^. Superimposing the structures of the RlmN-tRNA^Glu^ and CfrC105A/87-mer rRNA complexes reveals potential clashes between Cfr residues and nucleotides in the anticodon loop of tRNA^Glu^. Tyr252 clashes with C36; Lys318, and Thr325 clash with U34; His323 clashes with the phosphate group of U33; and Lys250 clashes with C32 (**Figure S7A**). These projected clashes may hinder the proper orientation and accommodation of the tRNA^Glu^ A37 nucleotide within the Cfr active site, supporting the observation that residues Lys250, Tyr252, and Thr325 are highly conserved. Furthermore, the superimposed structures indicate that the two substrates adopt similar conformations upon binding to their respective enzymes **(Figure S5**). For example, the anticodon stem of tRNA^Glu^ aligns with H90 of the rRNA; the acceptor stem corresponds to H91, and the D-arm of tRNA^Glu^ overlays with H92 of the 87-mer rRNA (**Figure S5**). Notably, the overall architecture of the 87-mer rRNA substrate adopts an L-shape that mimics the structure of tRNA (**Figures S4, S5**). Overlaying the H90 region of 23*S* rRNA from the 70*S E. coli* ribosome structure (PDB ID: 2AWB) with the Cfr-rRNA complex (**Figure S4C**) reveals distinct orientations of H91 and H92, suggesting that this region undergoes conformational changes during the late stages of ribosome maturation.

The Cfr C105A/87-mer rRNA structure also provides valuable clues for the observed preference shown by Cfr in methylating C^8^ over C^2^ of A2503. Close examination of the complex reveals that A2503 undergoes a 180° flip relative to the position of the A37 of tRNA^Glu^ in the RlmN-tRNA structure (PDB ID: 5HR7), which orients C^8^ of this nucleotide into a position that is suitable for methylation by Cfr. This observation might explain the conservation of amino acid residues within the active site radical-generating centers of the two enzymes (**Figure S6A**). However, the various residues highlighted in (**Figure S6B**) should dictate the correct orientation of C^8^ (in Cfr) or C^2^ (in RlmN) in the active site of the enzymes.

Recently, Fruci et al. reported an X-ray crystal structure of an inactive Cfr variant that contained 18 amino acid changes required for the protein to form crystals. The structure, determined to 2.50 Å resolution, was of the apo protein, lacking the [Fe_4_S_4_] cluster, SAM, and the RNA substrate. However, attempts were made to model these species into the structure^45^.

## Acknowledgements

We thank the members of the HHMI Janelia Research Campus Cryo-EM Shared Resources Facility and members of the Cryo-EM facility of the Penn State Huck Institutes of the Life Sciences for their helpful discussions and advice.

This work was supported by NIH (GM-122595, AI-180902), the Stephen and Patricia Benkovic Foundation, and the Eberly Family Distinguished Chair in Science to S.J.B. It was also supported by an NIH Award (S10OD026822) and Pennsylvania Department of Health Commonwealth Universal Research Enhancement (CURE) funds to S.L.H. S.J.B. is an investigator of the Howard Hughes Medical Institute.

## Data Deposition

The atomic coordinates and cryo-EM density maps will be available at the PDB and EMDB under accession codes listed in Table S1.

## List of Supplementary Materials

Table S1

Movie S2

SXYZ

SXYZ

