## Supporting Information for "Structural Basis for C^8^ methylation of 23S ribosomal RNA by Cfr"

**Supporting information includes:**

Materials and Methods

Table S1

Figures S1 to S7

**Materials and Methods**

*Materials.* Genes, codon-optimized for expression in *Escherichia coli*, were purchased from Invitrogen GeneArt Gene Synthesis Service through ThermoFisher Scientific (Waltham, MA). DNA oligonucleotide primers were obtained from Integrated DNA Technologies, Inc (Skokie, IL). Enzymes for RNA and DNA manipulation (Antarctic phosphatase, T7-RNAP, and restriction enzymes) were obtained from New England Biolabs, Inc. (Ipswich, MA). Lysozyme, DNase I, and P1 nuclease were purchased from Sigma-Aldrich (St. Louis, MO). Antibiotics, L-(+)-arabinose, dithiothreitol (DTT), and isopropyl-β-D-thiogalactopyranoside (IPTG) were purchased from Gold Biotechnology USA (St. Louis, MO). All other chemicals were purchased from Fisher Scientific at the highest purity available for biological applications. All solvents for liquid chromatography (LC) and mass spectrometry (MS) were of HPLC grade or higher. S-adenosylmethionine (SAM) was enzymatically synthesized following established protocols [1, 2].

*Methods. Cfr C105A variant expression and purification.* The expression and purification of the *Staphylococcus aureus* Cfr (UniProt ID A5HBL2) C105A variant with a C-terminal hexahistidine tag have already been described [3, 4].

*Synthesis and purification of rRNA substrates.* The 87-mer rRNA substrate was prepared by in vitro transcription. The DNA template, encompassing nucleotides 2496 to 2582 of *E. coli* 23S rRNA, was synthesized by PCR technology using the pKK3535 plasmid and the following primers: 5’-CGGAAATTAATACGACTCACTATAGGGCACCTCGA-3’ and 5′-mCmCAGCTCGCGTACCACTTTAAA-3′ (mC is 2-hydroxymethylcytosine) [5]. The resulting template was purified on a 1% agarose gel and used in a subsequent PCR to prepare additional template for in vitro transcription reactions. The 87-mer rRNA was synthesized in a T7 RNA polymerase (T7 RNA Pol) reaction containing 40 mM Tris-HCl, pH 8.4, 70 mM MgCl_2_, 20 mM DTT, 8 mM GTP, 4 mM CTP, 4 mM ATP, 4 mM UTP, 10 ng/μL 87-mer DNA template and 0.5 mg/ml of T7 RNA Pol in a final volume of 20 mL. The reaction was initiated by adding T7 RNA Pol. The transcription reaction was incubated at 37 °C for 2-4 h, and was quenched by adding EDTA to a final concentration of 110 mM. The precipitate was removed by centrifugation, and the reaction was concentrated to 5 mL using Amicon centrifugal filtration devices (Millipore Sigma, St. Louis, MO) with a 3 kDa molecular weight cut-off membrane. The concentrated RNA was introduced into a Coy anaerobic chamber and desalted using a Cytiva PD-10 column (Millipore Sigma) equilibrated in anaerobic, sterile water. The RNA was diluted to less than 100 μM in buffer containing 10 mM Tris-HCl, pH 8.0, and 50 mM NaCl, heated to 85 °C for 5 min, and slowly cooled to room temperature to allow for proper folding of the RNA secondary structure. The RNA was concentrated to ~1 mL and flash-frozen in liquid nitrogen. Its concentration was determined using an extinction coefficient of 1.01 μM^-1^ cm^-1^, calculated using OligoCalc (Thermo Fisher).

*Cfr C105A 87-mer Cross-link Preparation.* The Cfr/87-mer cross-link complex was prepared in a Coy anaerobic chamber. To form the cross-link, 400 μM Cfr C105A and 450 μM 87-mer substrate were combined in 20 mM Tris-HCl, pH 8.0, containing 20 mM MgCl_2_, and 2 mM SAM. Dithionite (2 mM) was added to initiate catalysis, and the reaction was incubated at room temperature for 2 h. The cross-link was purified by anion-exchange chromatography on a GE Healthcare HiPrep QFF column (Cytiva) interfaced with an AKTA purification system (Cytiva Life Sciences). The reaction mixture was diluted fivefold with Buffer A (50 mM Tris-HCl, pH 8.0, 50 mM NaCl, 5 mM DTT), loaded onto the column, and washed with 3 column volumes of Buffer A. Free protein, free RNA, and the cross-link complex were separated using a linear gradient from Buffer A to 50% Buffer B (50 mM Tris-HCl, pH 8.0, 2 M NaCl, 5 mM DTT) over 20 column volumes. The cross-link complex was collected and concentrated using a centrifugal filtration device with a 10 kDa molecular weight cut-off membrane. The concentration of the protein component was determined by Bradford analysis using a previously determined correction factor [3, 6].

*Cryo-EM Specimen Preparation and Data Collection*. The Cfr C105A/87-mer cross-link complex was exchanged into Buffer C (50 mM Tris-HCl, pH 8.0, 200 mM KCl, 2 mM MgCl_2_, 5 mM DTT, and 1% glycerol) to a final concentration of 1.3 mg/ml in untilted experiments or 0.8 mg/ml in tilted experiments. For the untilted datasets, Quantifoil R2/4 grids deposited with a single graphene monolayer fabricated by chemical vapor deposition (CVD) were used (Graphenea, San Sebastián, Spain). Using a graphene monolayer as an affinity substrate did not change Cfr’s particle orientation. The bias angle is orthogonal to the RNA overhang's long axis, which may have physically prevented the particles from rotating freely in an extremely thin ice environment. For tilted datasets, UltrAuFoil R1.2/1.3 grids (Quantifoil Micro Tools) were used instead to avoid the slightly elevated background signal from the graphene monolayer, which proved excessive for image signal from the very low molecular weight (~61.4 kDa) Cfr particles at a 30º stage tilt angle. The CVD graphene monolayer grids were gently glow-discharged for 15 s at 20 mA using a Pelco easiGlow system (Ted Pella Inc.), rendering the surface hydrophilic without noticeable damage to the graphene monolayer (**Figure S1 F**). The UltrAuFoil grids were glow-discharged for 30 s at 15 mA using the same PELCO easiGlow system. The sample was preincubated for 30 s on CVD graphene monolayer grids to ensure particle adsorption before blotting. The CVD graphene monolayer grids and UltrAuFoil grids were both blotted for 4 s at 0 blot force, 4 °C, and 100% relative humidity before vitrifyingvitrification in liquid ethane.

*Cryo-EM Data Collection.* Movies of Cfr particles embedded in vitreous ice were collected at close to liquid nitrogen temperature (77 K) using Titan Krios transmission electron microscopes (Thermo Fisher Scientific). The Cfr datasets were collected on the Janelia Krios1 equipped with a high brightness Field Emission Gun (X-FEG, Thermo Fisher Scientific) operated at 300 kV, a CETCOR spherical aberration (*C*s) corrector (Corrected Electron Optical Systems GmbH) with a educed Cs value of 0.01 mm, a Gatan BioContinuum HD energy filter operated with a 6 eV-wide energy slit, and a K3 direct electron detector (Gatan Inc). The K3 movies were recorded in super-resolution mode with correlative double-sampling (CDS) counting, and later binned 2×. Data were collected at a nominal magnification of 130,000×, corresponding to a calibrated magnification of 93,809×. A defocus range of -0.5 – -1.6 μm was applied. The total electron dose of each movie was 70 e^-^/Å^2^ fractionated over 70 frames. Semiautomated data collection was performed using SerialEM with customized scripts optimized for both data quality and throughput. The data collection parameters are in Table S1.

*Cryo-EM data processing.* Beam-induced motions of particles were corrected [7] using Unblur in *cis*TEM2 [8]. Contrast transfer function (CTF) parameters were estimated from sums of 3 movie frames using CTFFIND5 in *cis*TEM2 [8, 9]. The CTFFIND5 tilt search was used to estimate the CTF parameters for the dataset recorded at a 30º stage tilt angle. Particles were automatically picked *ab initio* using soft-edged disk templates generated internally in *cis*TEM2 [8, 10]. The picked particle images were boxed, extracted, and 2D-classified using *cis*TEM2 [8]. The initial 3D reconstructions were carried out *ab initio* [11], followed by initial 3D refinements using *cis*TEM2 with refinement of the beam tilt and per-particle CTF parameters [8, 11]. 3D classifications and final 3D refinement were carried out using Relion5 with beta Blush regularization (**Figure S1A**) [12, 13]. A 3D variability analysis (3DVA) was performed using cryoSPARC v4 to visualize the extent of conformational heterogeneity present [14]. The number of movies and particles used in the final reconstructions is listed in **Table S1**.

*Model building.* An alpha-fold predicted structural model of Cfr [15] was used as the initial model and manually rebuilt using COOT [16, 17]. All protein models were real-space-refined using PHENIX [18] and evaluated using COOT, ISOLDE, and the MolProbity server [19, 20]. The cryo-EM maps were deposited in the Electron Microscopy Databank (EMDB), and the coordinates of the atomic model were deposited in the Protein Data Bank (PDB) [21, 22]. Figures were generated using PyMOL, WARP, Chimera, ChimeraX, and Adobe Illustrator [23, 24].

**Table S1 | Cryo-EM data collection, refinement and validation statistics.**

|  | Cfr in complex with 87-mer 23*S* rRNA  (EMDB-xxxx)  (PDB xxxx) |
| --- | --- |
| **Data collection and processing** |  |
| Magnification | 130,000 |
| Voltage (kV) | 300 |
| Electron exposure (e^–^/Å^2^) | 70 |
| Defocus range (μm) | -0.5 to -1.6 |
| Pixel size (Å) | 0.533 |
| Symmetry imposed | C1 |
| Initial particle images (no.) | 5,314,998 |
| Final particle images (no.) | 74,472 |
| Map resolution (Å)  FSC threshold | 3.0  0.143 |
| Map sharpening *B* factor (Å^2^) | -98.54 |
| **Refinement** |  |
| Initial model used (PDB code) | - |
| Model composition  Non-hydrogen atoms  Protein residues  Nucleic acid residues  Ligands  Refinement (Phenix)  Map correlation coefficient  (whole unit cell)  Map correlation coefficient (around atoms)  R.m.s. deviations  Bond lengths (Å)  Bond angles (°) | 4,134  343  63  3  0.82  0.78  0.004  0.764 |
| Validation  MolProbity score  Clashscore  Poor rotamers (%) | 1.69  10.11  0.00 |
| Ramachandran plot  Favored (%)  Allowed (%)  Disallowed (%) | 97.04  2.96  0.00 |

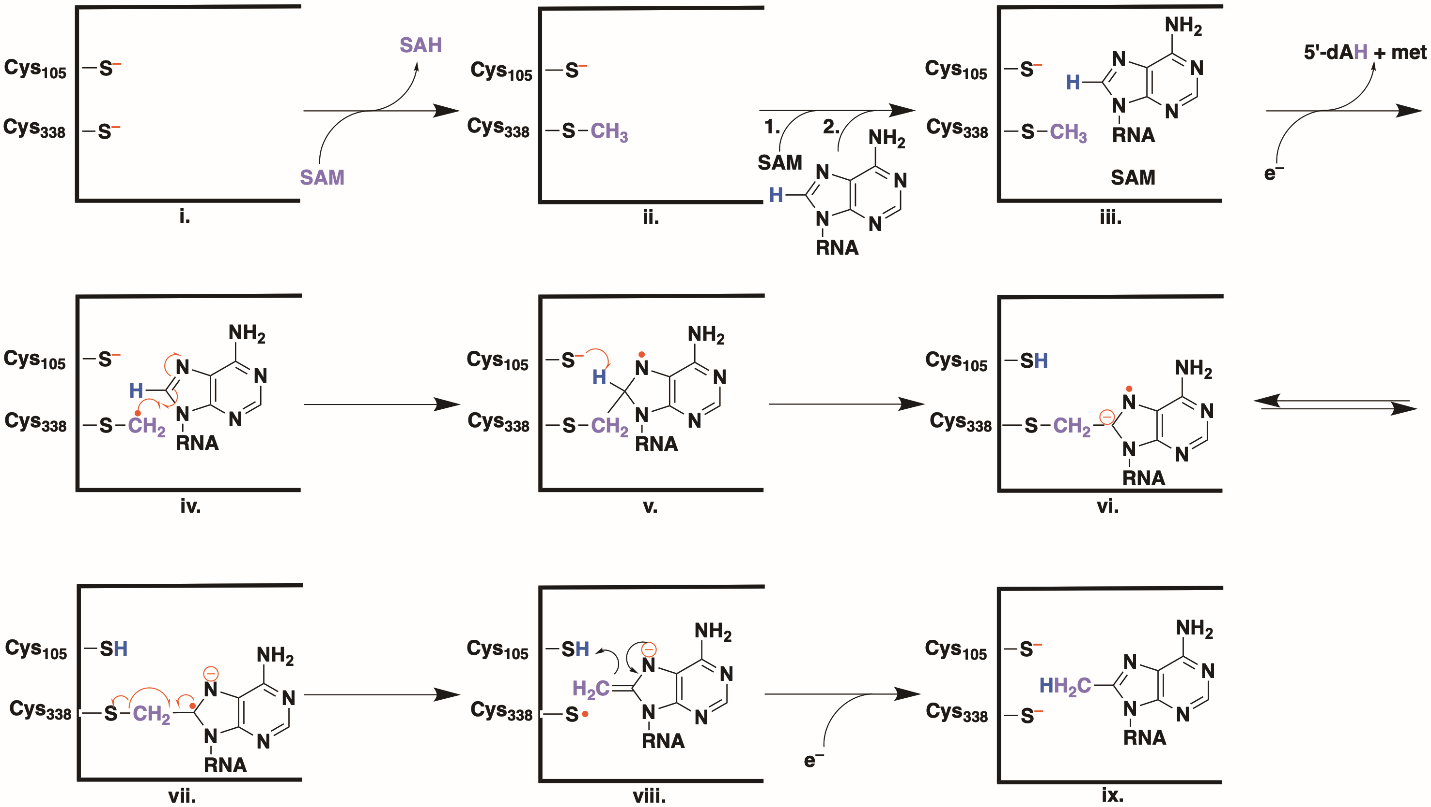

**Figure S1**. Proposed Cfr reaction mechanism

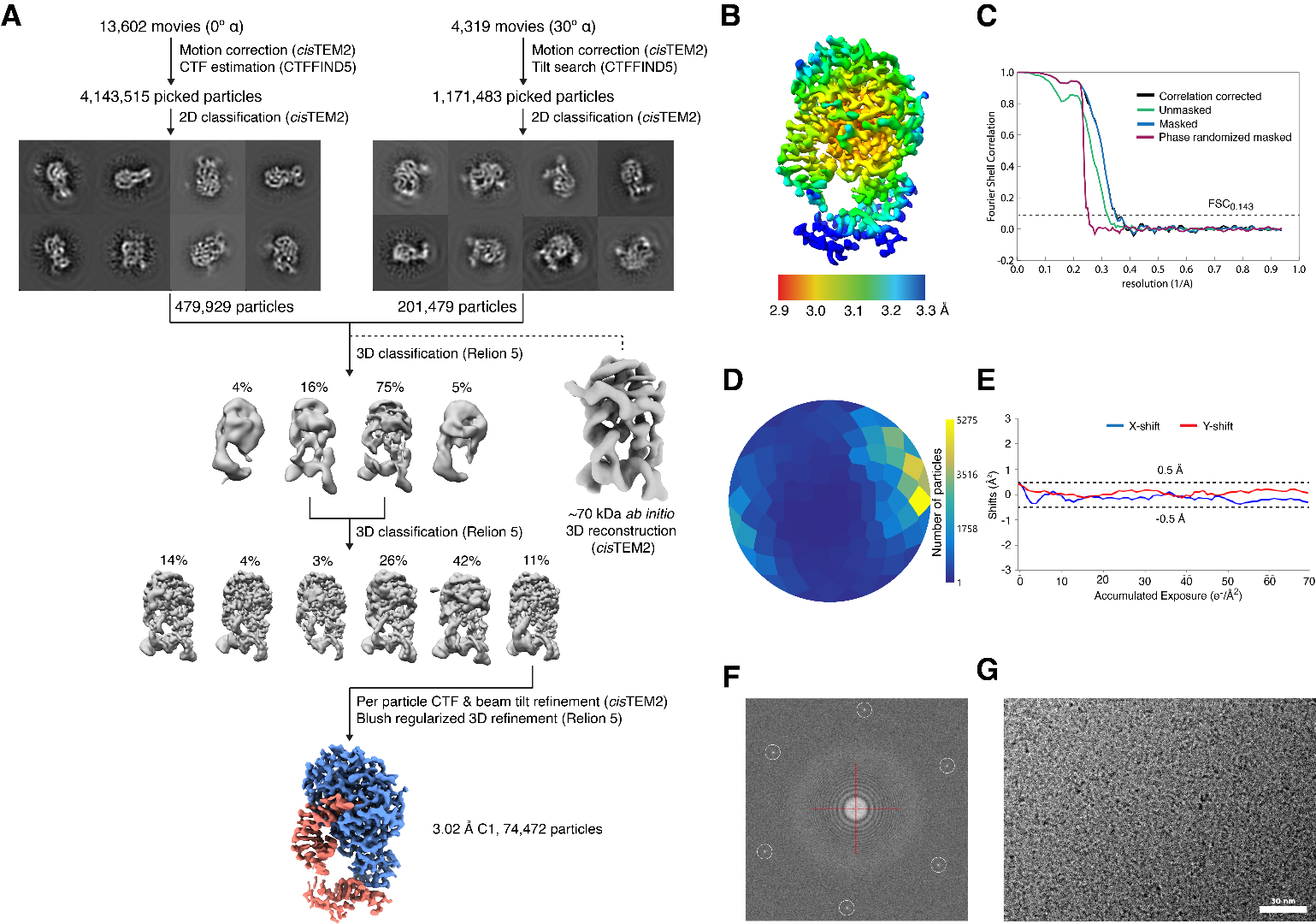

**Figure S2. (A)** Cryo-EM data processing. **(B)** Local resolution map of the CfrC105A/87-mer rRNA complex. The surface-rendered map of the complex structure was colored for local resolution according to the key, with most regions ranging from 2.9 to 3.3 Å resolution. (C) Fourier shell correlation (FSC) plots. **(D)** The angular distribution of particle projections calculated in RELION and WARP [13, 24]. **(E)** CVD graphene monolayer suppressed beam-induced motion to a sub-0.5-1 Å level on 2.0 μm diameter holes. (A) **(F)** Fast Fourier transform of CfrC105A/87-mer rRNA cryo-EM micrograph with graphene monolayer reciprocal lattice (in circles). **(G)** Cryo-EM micrograph of CfrC105A/87-mer rRNA complex.

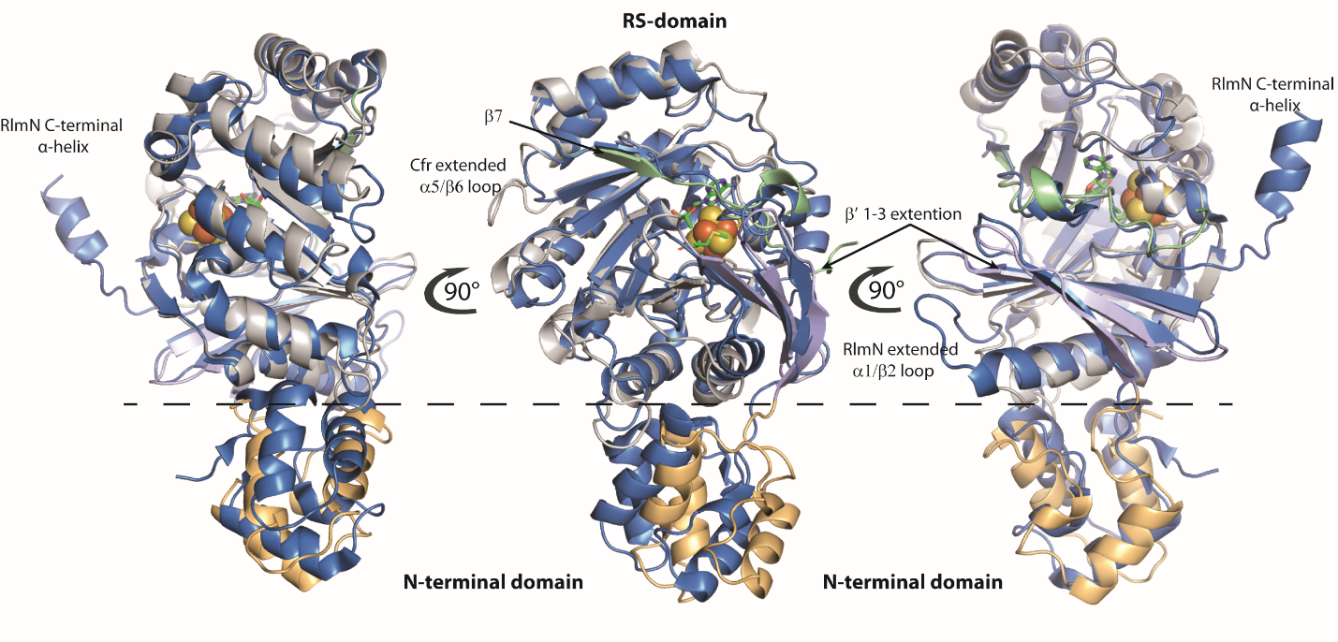

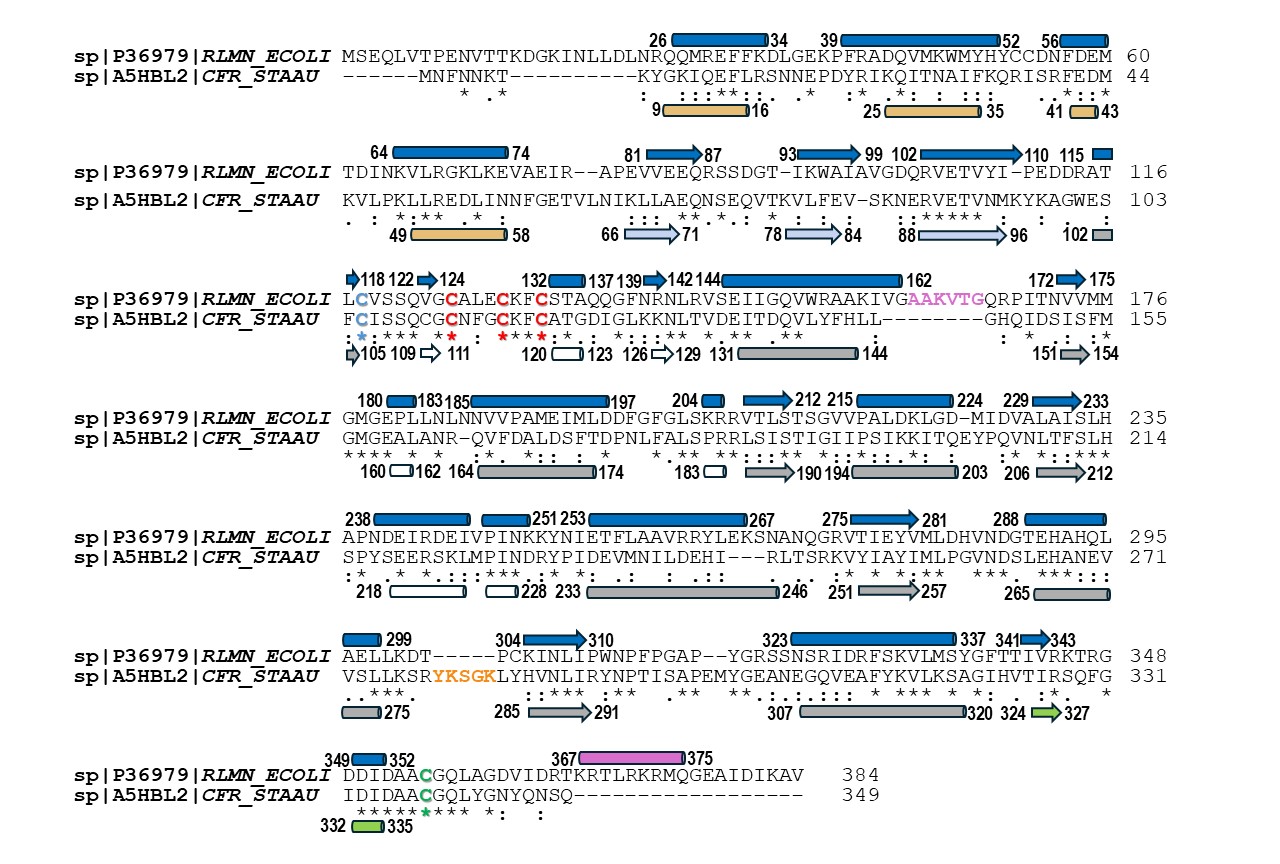

**A**

**B**

**Figure S3. (A)** A cartoon representation of the superimposed structures of EcRlmN (PDB ID: 5HR7, blue color) and CfrC105A (N-terminal domain (tan); β’1-3 extension (light blue); radical SAM domain (light grey); C-terminal (green)). **(B)** Sequence alignment of *E. coli* RlmN and Cfr with secondary-structure annotations. Secondary structure elements are depicted as cylinders (α-helices) and arrows (β-strands). The color code matches that used in panel A, except that the TIM-barrel fold of the RS domain of Cfr is shown in grey and the RS domain extensions are shown in white. The Cys residues that coordinate the [Fe_4_S_4_] cluster are shown in red; the Cys residue changed to Ala for forming the crosslinked protein–RNA complex is shown in blue; and the Cys residue transiently methylated during catalysis is shown in green. The loop sequence and α-helix shown in pink are observed only in RlmN, while the orange-colored loop sequence is observed only in Cfr.

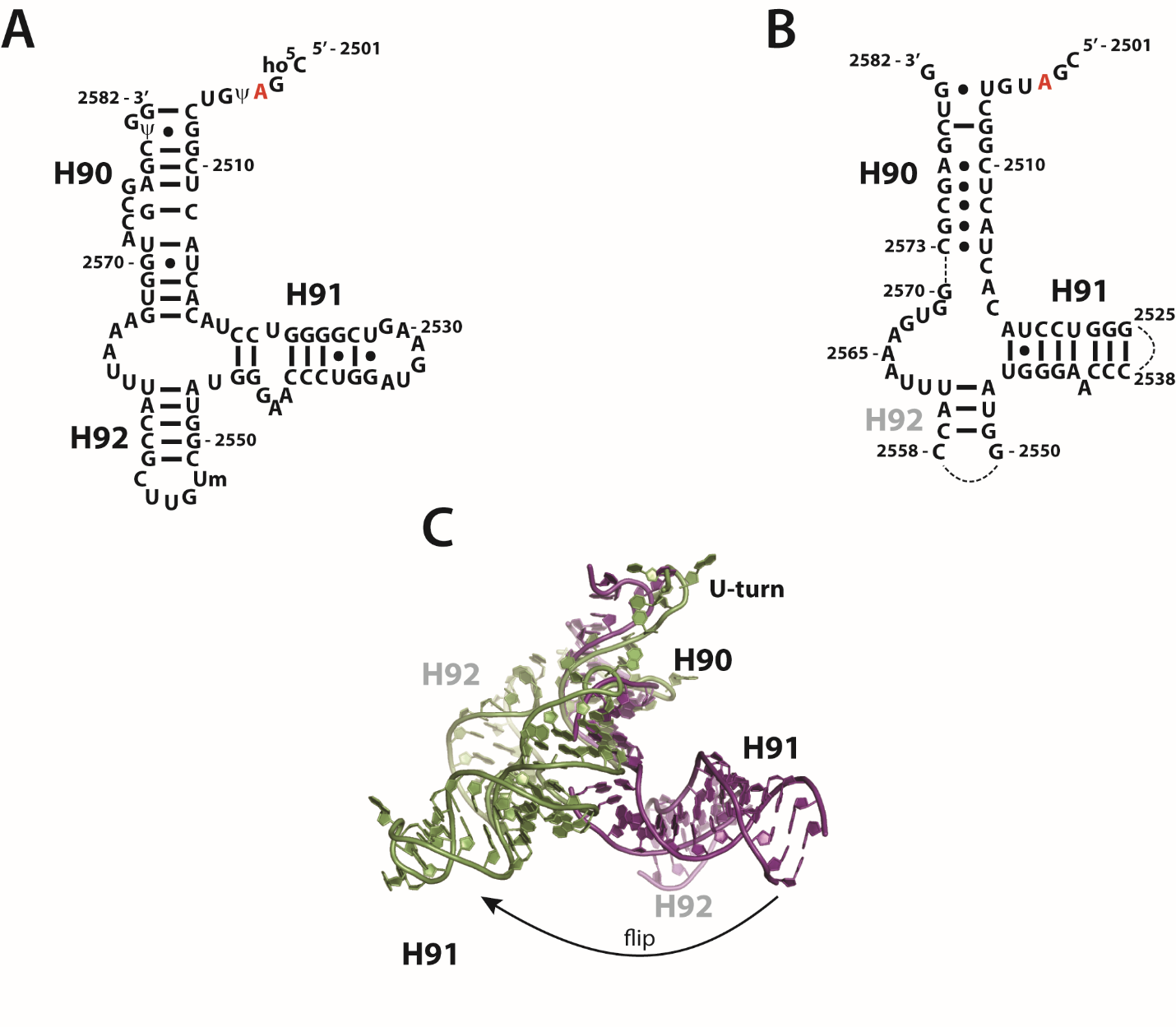

**Figure S4. (A)** Secondary structure of 23S rRNA fragment corresponding to nucleotides 2501 to 2582 of native rRNA. (**B**) Secondary structure of the 87-mer fragment from the structure of the Cfr C105A/87-mer rRNA complex. (**C**) Superimposed structures of the native 23S rRNA fragment (nucleotides 2501 to 2582) from the ribosome (PDB ID: 2AWB, green) and the 87-mer RNA fragment from the crosslinked structure of Cfr C105A/87-mer (purple color).

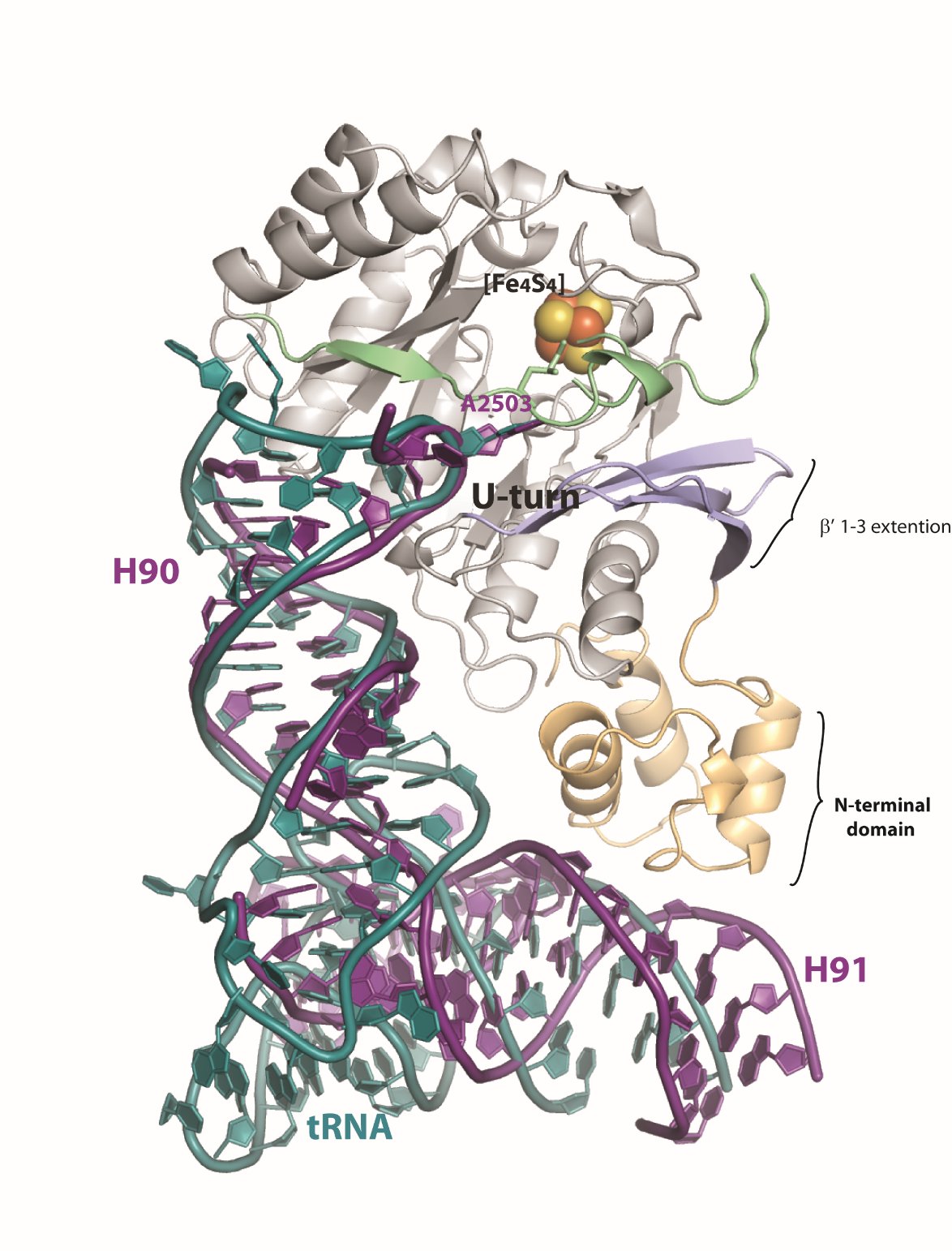

**Figure S5.** Superimposed structures of the EcRlmN C118A - tRNA^Glu^ (PDB ID: 5HR7) and the Cfr C105A/87-mer rRNA complexes showing similar L-shape structures of the RNA substrates. Color code: N-terminal domain (tan); β’1-3 extension (light blue); radical SAM domain (light grey); C-terminal region (light green); rRNA fragment spanning nucleotides 2501 to 2582 (purple); tRNA^Glu^ (green).

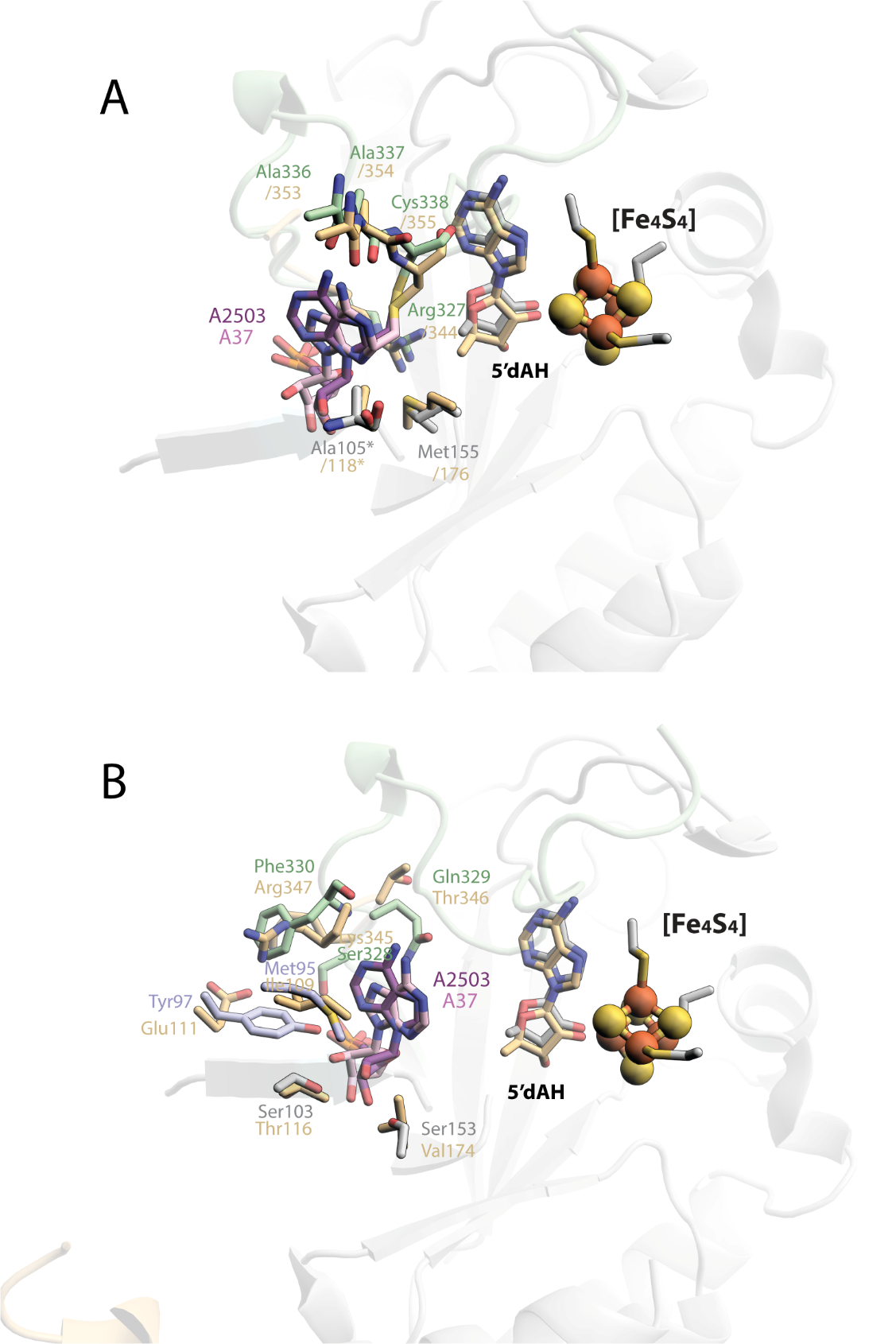

**Figure S6.** Superimposed active sites of the E. coli RlmN C118A - tRNA^Glu^ complex (PDB ID: 5HR7) with the CryoEM structure of Cfr C105A/87-mer rRNA, showing (A) identical residues, and (B) nonidentical residues in the active sites of E. coli RlmN and Cfr. Color code: E. coli RlmN residues (tan); Cfr residues from β’1-3 extension (light blue), from radical SAM domain (light grey), from C-terminal region (light green); A2503 of rRNA (purple); A37 of tRNA^Glu^ (pink).

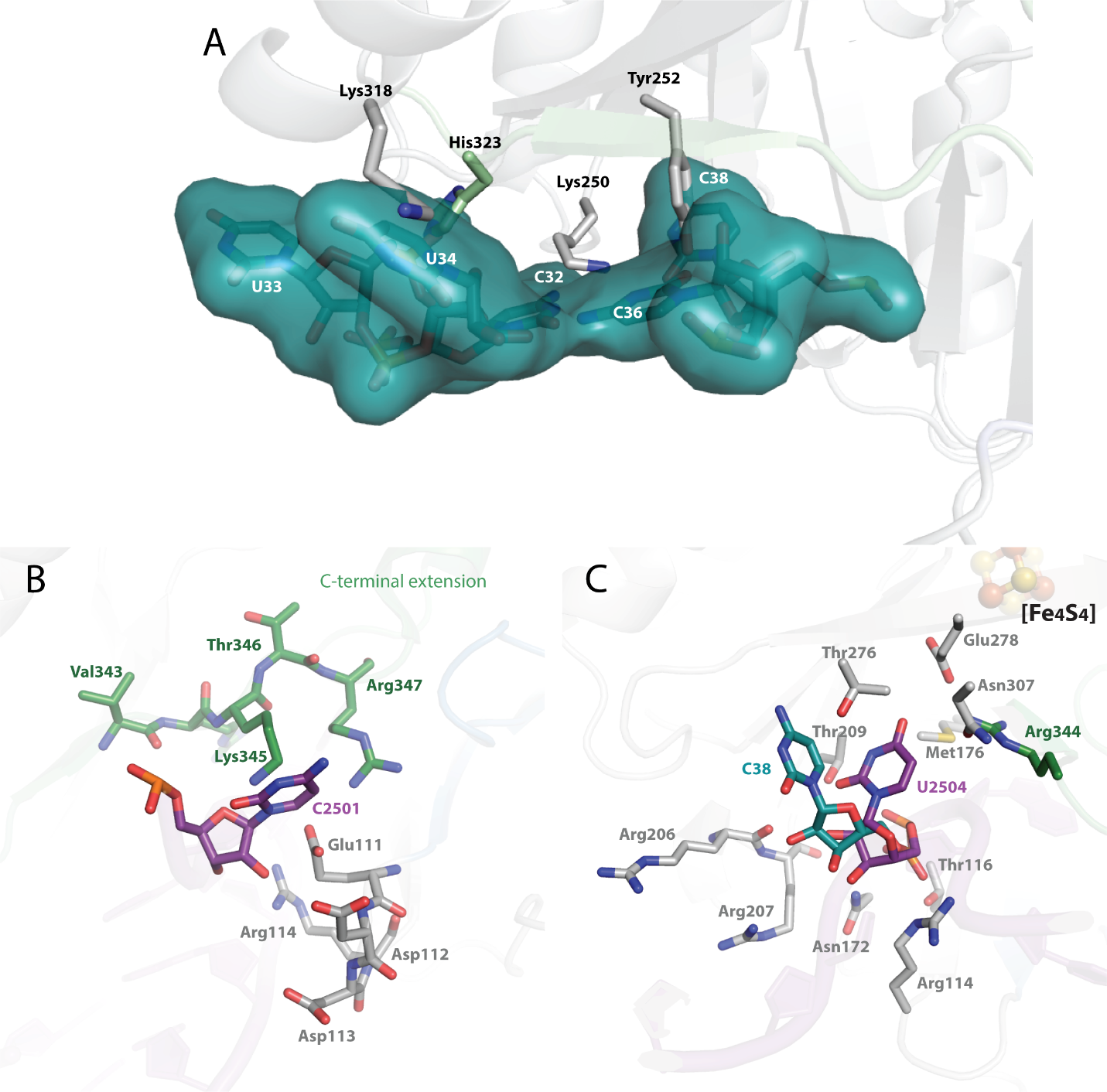

**Figure S7.** An overlay of the Cfr C105A structure with that of tRNA^Glu^ from the EcRlmN C118A - tRNA complex (PDB ID: 5HR7) **(A)** shows clashes between residues of Cfr and the tRNA anticodon loop. Color code: tRNA anticodon loop (turquoise sticks and surface in panel A); Cfr residues from the β’1-3 extension (light blue), from the radical SAM domain (light grey), from the C-terminal region (light green) Predicted positions of cytosine 2501 (C2501) (**B**) and uridine 2504 (U2504) (**C**) of 23S rRNA in the active site of RlmN. Color code: rRNA (purple); RlmN residues from the radical SAM domain (grey), from the C-terminal region (green); C38 of the tRNA anticodon stem loop is shown in turquoise.
